# Cortical and thalamic connectivity of temporal visual cortical areas 20a and 20b of the domestic ferret (*Mustela putorius furo*)

**DOI:** 10.1101/492728

**Authors:** Dell Leigh-Anne, Giorgio M. Innocenti, Claus C. Hilgetag, Paul R. Manger

## Abstract

The present study describes the ipsilateral and contralateral cortico-cortical and corticothalamic connectivity of the temporal visual areas **20a** and **20b** in the ferret using standard anatomical tract-tracing methods. The two temporal visual areas are strongly interconnected, but area **20a** is primarily connected to the occipital visual areas, whereas area **20b** maintains more widespread connections with the occipital, parietal and suprasylvian visual areas and the secondary auditory cortex. The callosal connectivity, although homotopic, consists mainly of very weak anterograde labelling which was more widespread in area **20a** than area **20b**. Although areas **20a** and **20b** are well connected with the visual dorsal thalamus, the injection into area **20a** resulted in more anterograde label, whereas more retrograde label was observed in the visual thalamus following the injection into area **20b**. Most interestingly, comparisons to previous connectional studies of cat areas **20a** and **20b** reveal a common pattern of connectivity of the temporal visual cortex in carnivores, where the posterior parietal cortex and the central temporal region (**PMLS**) provide network points required for dorsal and ventral stream interaction enroute to integration in the prefrontal cortex. This pattern of network connectivity is not dissimilar to that observed in primates, which highlights the ferret as a useful animal model to understand visual sensory integration between the dorsal and ventral streams. This data will contribute to the Ferretome (www.ferretome.org) to facilitate cross species analysis of brain connectomes and wiring principles of the brain.

## Introduction

The size of the temporal region of the mammalian cerebral cortex varies greatly amongst species, with this region being expanded and seemingly specialized for the processing of complex visual perceptions in primates (for review see Leopold, Mitchell & Freiwald, 2017). Due to the expansion of the temporal cortex in primates, and the obvious relation to understanding object recognition capacities in humans, the study of temporal cortex in non-primate mammals has not be a priority. Indeed, in many commonly studied species the temporal cortex has not been investigated and defined, due to its small size and the difficulty of access to this region due to its location within the skull. Despite this obstacle, even in species with unexpanded temporal cortices relative to primates, such as sheep, goats and dogs, face-recognition neurons and the ability to recognize faces have been described (e.g Kendrick, da Costa, Leigh, Hinton, & Pierce, 2001; Tate, Fischer, Leigh & Kendrick, 2006). In primates it has been well-established that while the temporal visual areas are connected with other adjacent visual areas, they also possess two sets of long range connections, to the posterior parietal cortex and to the orbitofrontal cortex (e.g. Webster, Bachevalier, & Ungerleider, 1994). It is thought that these long range connections are involved in the social aspects of vision as well as visually guided behaviour in relation to identified objects (Leopold et al., 2017).

Thus, in order to understand the baseline from which the complex temporal visual cortex of primates, with its associated range of visual abilities, evolved, it is important to understand the organization of these regions in other species, such as the ferret, who possess a relatively unexpanded temporal cortex. In both the cat and ferret a cluster of three cortical visual areas (**20a, 20b, PS**), located lateral to the occipital visual areas (**17, 18, 19, 21**) and medial to the perirhinal cortex, have been defined as temporal visual areas (Tusa & Palmer, 1980; Updyke, 1986; Manger, Nakamura, Valentiniene, & Innocenti, 2004; Homman-Ludiye, Manger, & Bourne, 2010), despite some differences in the precise organization of the retinotopic maps in each of these areas between ferret and cat (Manger et al., 2004). Studies of intrahemispheric connectivity of these temporal regions in the cat have revealed connections with occipital (Symonds & Rosenquist, 1979; Tusa & Palmer, 1980), suprasylvian (Heath & Jones, 1971; Symonds & Rosenquist, 1979), and parietal visual areas (Cavada & Reinoso-Suárez, 1983; Avendaño, Rausell, Perez-Aguilar, & Isorna, 1988), as well as with frontal cortical areas (Cavada & Reinoso-Suárez, 1981, 1983; Olson & Jeffers, 1987); however, it is difficult to propose direct cortical area homologs and compare the connectional data of the cat with species with a more complexly organized temporal cortex such as the macaque monkey (see Payne, 1993; Manger et al., 2004). It is likely to be more beneficial to our understanding of cortical processing to consider different regions, such as occipital, temporal, parietal, as a whole and determine whether global underlying connectivity patterns are present across species. The global interhemispheric connectivity of the ferret temporal cortical region has been studied (Manger et al., 2004), and, in addition, these ferret temporal visual areas appear to be strongly connected to the higher order auditory areas of posterior ectosylvian gyrus (Bizley & King, 2009); however, no studies have examined the specific cortical and thalamic connectivity of the individual temporal visual areas of the ferret.

Here we analyze and describe cortico-cortical and cortico-thalamic connectivity of ferret temporal visual areas **20a** and **20b**. We anticipate that these studies may provide clues about the baseline mammalian substrate for multisensory processing and integration with the temporal cortex, and may provide insights into the anatomical interaction and connectivity of the dorsal and ventral processing streams required for the execution of complex visual and auditory behaviours.

## Material and Methods

### Surgical procedure and tracer injections

Four adult female ferrets (*Mustela putorius*), weighing between 600g and 1000g, were used in this current study (two injection sites per cortical area). The experiments were conducted according to the Swedish and European Community guidelines for the care and use of animals in scientific experiments. All animals were initially anesthetized with i.m. doses of ketamine hydrochloride (Ketalar, 10mg/kg) and medetomidin hydrochloride (Domitor, 0.08mg/kg), supplemented with atropine sulphate (0.15mg/kg) and placed in a stereotaxic frame. A mixture of 1% isoflurane in a 1:1 nitrous oxide and oxygen mixture was delivered through a mask while the animal maintained its own respiration. Anesthetic level was monitored using the eye blink and withdrawal reflexes, in combination with heart rate measurement. The temporal cortex was exposed under aseptic conditions and in each animal numerous (but less than 20) electrophysiological recordings were taken to ensure correct placement of the tracer within a specific cortical area (Manger et al., 2004). Approximately 500 nl of tracer (biotinylated dextran amine, BDA 10 k, 5% in 0.1 M phosphate buffer; Molecular Probes) was delivered at each injection site using a Hamilton microsyringe (Figs. 1, 2a, 2b). After the completion of the injections, a soft contact lens was cut to fit over the exposed cortex, while the retracted dura mater was pulled over the contact lens and the excised portion of bone repositioned and held in place with dental acrylic. The temporal muscle was reattached using surgical glue and the midline incision of the skin sutured. Antibiotics were administered to all cases (Terramycin, 40 mg/kg, daily for 5 days) and these animals were given a 2-week recovery period to allow for tracer transport. At the end of this period, the animals were euthanized with a lethal dose of sodium pentobarbital (80 mg/kg, i.p.) and perfused intracardially, initially with a rinse of 0.9% saline (4°C, 500ml/kg), followed by fixation with 4% paraformaldehyde in 0.1M phosphate buffer (4°C, 1000 ml/kg).

**Figure 1:**
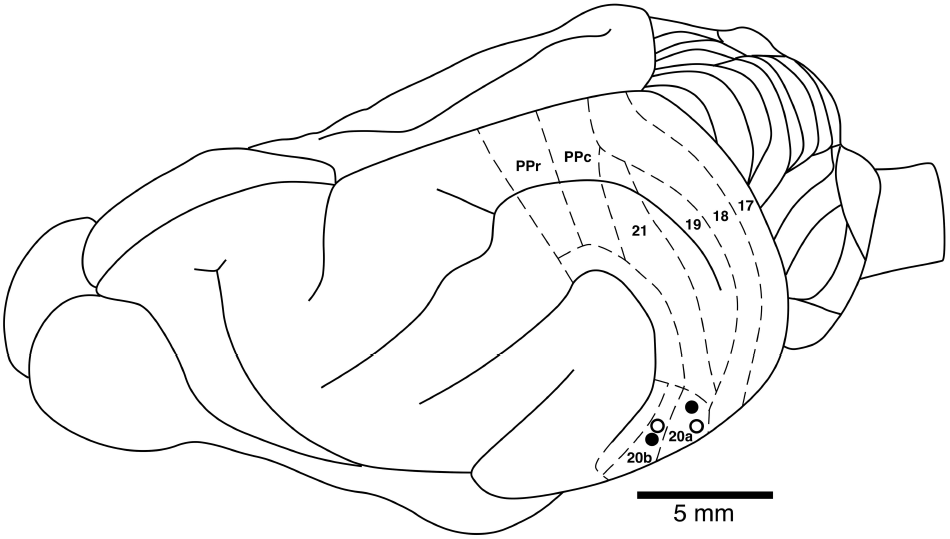
Locations of the temporal visual areas injection sites analyzed in the current study in relation to many of the known boundaries of visual cortical areas in a dorsolateral view of the ferret brain. Closed circles represent the injections sites where the brain was sectioned in a coronal plane. Open circles represent the injections sites where the cerebral cortex was manually semi-flattened for analysis. See list for abbreviations.

### Sectioning and staining procedures

The brains were removed from the skull and post-fixed overnight in 4% paraformaldehyde in 0.1M phosphate buffer and then transferred to a 30% sucrose solution in 0.1M phosphate buffer (4°C) and allowed to equilibrate. The brains were either: (1) frozen in dry ice and sectioned at 50 μm on a freezing microtome in a coronal plane (2 cases, one for each of the temporal cortical areas 20a and 20b) for a one in four series for Nissl (cresyl violet), myelin (Gallyas, 1979), cytochrome oxidase (Carroll & Wong-Riley, 1984) and BDA; or (2) cryoprotected and the cerebral cortex dissected away from the remainder of the brain and the dorsolateral surface semi-flattened (2 cases, one for each of the temporal cortical areas 20a and 20b) between two glass slides, frozen onto the cold microtome stage and sectioned parallel to the semi-flattened surface at 50 μm for a one in two series for BDA and cytochrome oxidase. For BDA tracer visualization, the sections were incubated in 0.5% bovine serum albumin in 0.05M Tris buffer for 1h, followed by incubation in an avidin-HRP solution for 3h. A 10 min pre-incubation in 0.2% NiNH_4_SO_4_ preceded the addition of H_2_O_2_ (200 μl/l) to this solution, at which time the sections were monitored visually for the reaction product. To stop the reaction the sections were placed in 0.05 M Tris buffer. All sections were mounted on 0.5% gelatine coated slides, dehydrated in a graded series of alcohols, cleared in xylene and coverslipped with Depex mounting medium. All injection sites resulted in robust anterograde and retrograde transport of tracer in both the cerebral cortex (Fig. 2c-f) and the visual portions of the dorsal thalamus (Fig. 3).

**Figure 2:**
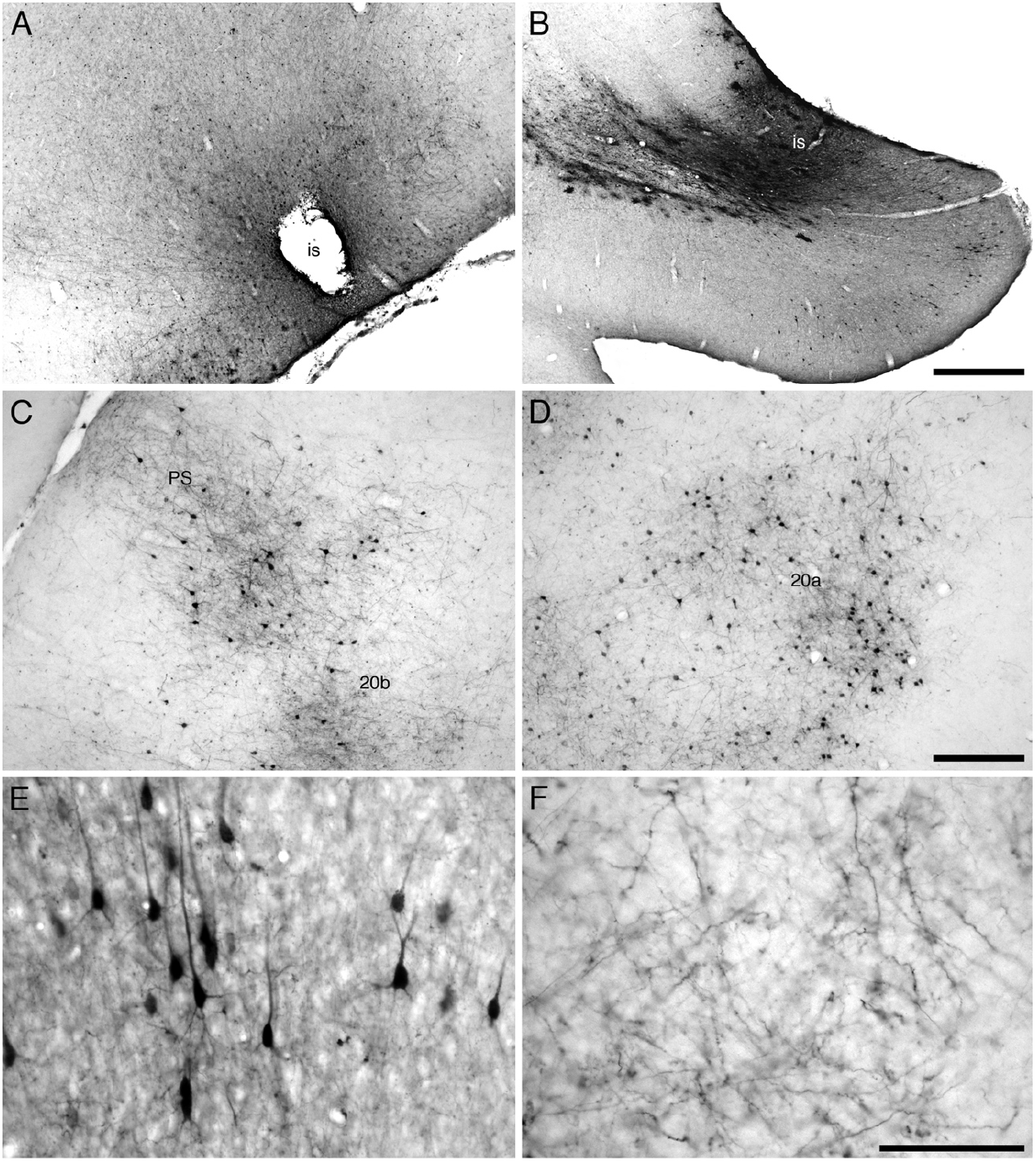
Photomicrographs showing examples of injection sites (**a, b**) and labelled cells (**c-f**) that were analyzed in the current study. (**a**) Injection site (**is**) in area **20a** of a semi-flattened cerebral cortex showing the spread of tracer around the injection site, indicating the areal specificity of the injections made in the current study. (**b**) Injection site (**is**) in area **20b** from a coronal section through the cerebral cortex, showing the spread of tracer and the limitation of the injection site to the cerebral cortex. Labelled cells and terminals in areas **PS** and **20b** (**c**) and **20a** (**d**) following transport from the injection sites in area **20a** (**c**) and area **20b** (**d**). (**e**) High magnification image of a retrogradely labelled cell in area **20b** following transport from the injection site depicted in **b**. (**f**) High magnification image of anterogradely labelled axons in area **19** following transport from the injection site depicted in **a**. Scale bar in **b** = 500 μm and applies to **a** and **b**. Scale bar in **d** = 250 μm and applies to **c** and **d**. Scale bar in **f** = 100 μm and applies to e and f. In images **a, c, d**, and **f**, the midline of the brain is to the top of the image and rostral to the left. In image **b**, medial is to the left and dorsal to the top. In image e the pial surface has been rotated to the top of the image.

**Figure 3:**
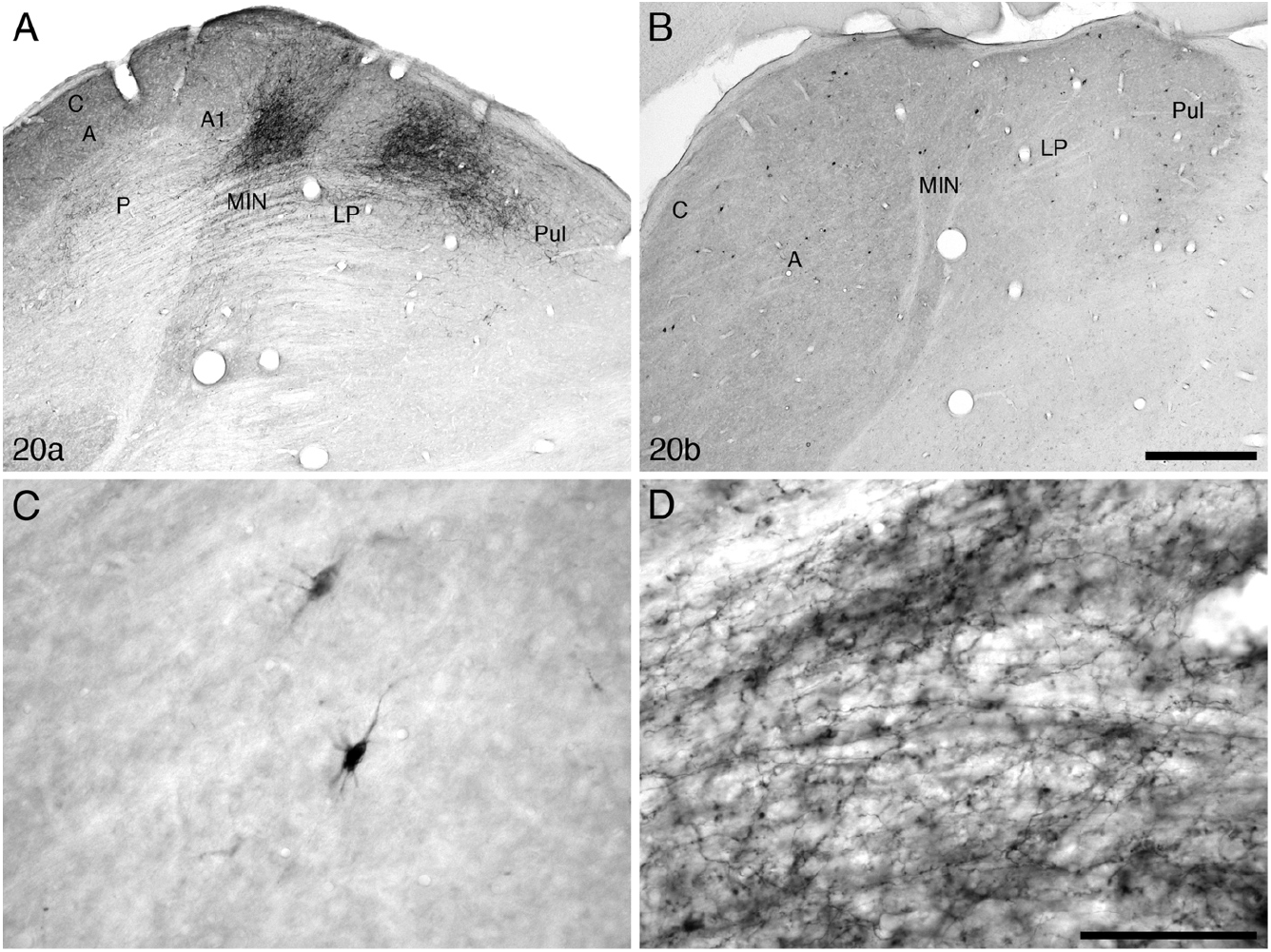
Photomicrographs showing examples of retrogradely labelled cells and anterogradely labelled axons in the visual thalamus of the ferret following injections into the temporal visual areas **20a** (**a**), and **20b** (**b**). (**c**) High magnification image of retrogradely labelled cells in the lateral geniculate nucleus following transport from the injection site in area **20b**. (**d**) High magnification image of anterogradely labelled axons in the lateral posterior nucleus following transport from the injection site in area **20a**. Scale bar in **b** = 500 μm and applies to **a** and **b**. Scale bar in **d** = 100 μm and applies to **c** and **d**. In all images dorsal is to the top and medial to the right. See list for abbreviations.

### Qualitative and quantitative analysis

For qualitative analysis, the stained sections were examined under low and high power magnification using a light microscope to determine in which sections through the cortex labelled cell bodies and terminals were present. Under low power stereomicroscopy using the flattened sections, the edges of each section were drawn with the aid of a camera lucida, and the location of the injection site marked. Areal borders were delineated and drawn using the cytochrome oxidase stained sections. The sections reacted for BDA were then matched to these drawings and the locations of the individual retrogradely labelled cells plotted and regions of anterogradely labelled axonal terminals demarcated. The drawings were scanned and redrawn using a Canvas X Pro 16 drawing program (ACD Systems International Inc., USA). Digital photomicrographs were captured using a Zeiss Axioskop and the Axiovision software. No pixilation adjustments or manipulation of the captured images were undertaken, except for contrast and brightness adjustment using Adobe Photoshop 7.

To quantify the retrograde BDA labelling per injection site, and control for variance in the size of the injection, a fraction of labelled neurons (N%, Table 1) was calculated using the formula:

> N% = Σ Projection neurons identified within a region / Σ Total projection neurons identified across both hemispheres × 100

Where cell bodies could be clearly identified, neurons within the injection halo were counted. Furthermore, cell bodies located on boundary lines between areas were only accounted for in the region where the most overlap was present.

**Table 1:** Average fraction of retrogradely labelled neurons (N%), in the various visual cortical areas of the ipsilateral and contralateral hemispheres of the ferret following injections of biotinylated dextran amine (BDA) into the temporal visual areas **20a** and **20b**. See list for abbreviations. **IS** – injection site.

|  | Occipital |  |  |  | Temporal |  |  |  | Auditory |  | Suprasylvian |  |  |  |  |  | Parietal |  | Somatomotor |  |  |
| --- | --- | --- | --- | --- | --- | --- | --- | --- | --- | --- | --- | --- | --- | --- | --- | --- | --- | --- | --- | --- | --- |
| IS | 17 | 18 | 19 | 21 | 20a | 20b | PS | AEV | A1° | A2° | PMLS | PLLS | AMLS | ALLS | VLS | DLS | PPc | PPr | SS | Mot | PreF |
| **Ipsilateral connectivity** |  |  |  |  |  |  |  |  |  |  |  |  |  |  |  |  |  |  |  |  |  |
| **20a** | 2.39 | 21.21 | 12.51 | 19.26 | 16.41 | 15.10 | 2.07 | 0.00 | 0.15 | 1.51 | 2.24 | 0.12 | 0.44 | 0.15 | 2.33 | 0.32 | 0.06 | 0.06 | 0.06 | 0.03 | 0.00 |
| **20b** | 0.00 | 6.95 | 2.47 | 13.40 | 5.95 | 21.42 | 9.93 | 2.24 | 0.11 | 14.93 | 7.01 | 1.96 | 0.17 | 2.08 | 5.83 | 0.56 | 4.54 | 1.79 | 0.50 | 0.34 | 0.22 |
| **Contralateral connectivity** |  |  |  |  |  |  |  |  |  |  |  |  |  |  |  |  |  |  |  |  |  |
| **20a** | 0.35 | 0.44 | 0.20 | 1.11 | 1.25 | 0.06 | 0.09 | 0.00 | 0.00 | 0.06 | 0.00 | 0.00 | 0.00 | 0.00 | 0.00 | 0.00 | 0.00 | 0.00 | 0.06 | 0.00 | 0.00 |
| **20b** | 0.00 | 0.11 | 0.00 | 0.11 | 0.62 | 0.06 | 0.17 | 0.00 | 0.00 | 0.39 | 0.00 | 0.00 | 0.00 | 0.00 | 0.17 | 0.00 | 0.28 | 0.17 | 0.00 | 0.00 | 0.00 |

## Abbreviations

3a: rostral, or proprioceptive, somatosensory area
17: primary visual cortex
18: second visual cortical area
19: third visual cortical area
20a: temporal visual area a
20b: temporal visual area b
21: fourth visual cortical area
A: A lamina of the lateral geniculate nucleus
A1: A1 lamina of the lateral geniculate nucleus
AI: primary auditory cortex
AAF: anterior auditory field
ADF: anterior dorsal auditory field
AEV: anterior ectosylvian visual area
AMLS: anteromedial lateral suprasylvian visual area
ALLS: anterolateral lateral suprasylvian visual area
AVF: anterior ventral auditory field
C: C lamina of the lateral geniculate nucleus
DLS: dorsal lateral suprasylvian visual cortical area
is: injection site
LP: lateral posterior nucleus of the dorsal thalamus
M1: primary motor cortex
MIN: medial intralaminar nucleus of the lateral geniculate nucleus
OB: olfactory bulb
OT: olfactory tract
P: perigeniculate lamina of the lateral geniculate nucleus
PIR: piriform cortex
PMLS: posteromedial lateral suprasylvian visual area
PLLS: posterolateral lateral suprasylvian visual area
PPc: posterior parietal caudal cortical area
PPF: posterior pseudosylvian field
PPr: posterior parietal rostral cortical area
PreF: prefrontal cortical region
PreM: premotor cortical region
PS: posterior suprasylvian visual cortical area
PSF: posterior suprasylvian auditory field
Pul: pulvinar nucleus of the dorsal thalamus
SI: primary somatosensory area
SII/PV: second somatosensory area/parietoventral somatosensory area
SIII: third somatosensory area
SMA?: potential supplementary motor area
SomS: somatosensory cortex
Vb: ventrobasal complex of the dorsal thalamus
VLS: ventral lateral suprasylvian visual area
VP: ventral posterior ectosylvian region.

## Results

In the current study, BDA tracer injections were made into the temporal visual cortical areas **20a** and **20b** of the ferret, in order to assess the distribution of both anterograde and retrograde cortico-cortical and cortico-thalamic connections (Figs. 1 – 3). In all cases there was more anterograde and retrograde label present in the hemisphere ipsilateral to the injection site than in the contralateral hemisphere, and this label was present in patches or clusters. Visual inspection revealed that the injection into **20a** produced denser anterograde and retrograde ipsilateral labeling than that of **20b**, but the ipsilateral connections from the **20b** injections were substantially more widespread across the cortex than those from the **20a** injections. Quantification of the fraction of labeled neurons (N%) showed that more than 96% of labeled cell bodies were found in the hemisphere ipsilateral to the injection site, with a substantial proportion of the labeled neurons being found within the cortical area injected (Table 1, Fig. 4). Furthermore, area **20a** was primarily connected with the occipital and suprasylvian visual areas, whereas area **20b** was connected with suprasylvian, parietal, auditory and pre-motor areas. Ipsilateral reciprocal connectivity was observed throughout the visual nuclei of the dorsal thalamus, and no contralateral connections to the visual thalamus were observed. Substantially more retrograde labeling was identified in the visual dorsal thalamus following the tracer injection into area **20b** compared with **20a**, but the **20a** injection resulted in more anterograde labeling in the visual dorsal thalamus than the **20b** injection.

**Figure 4:**
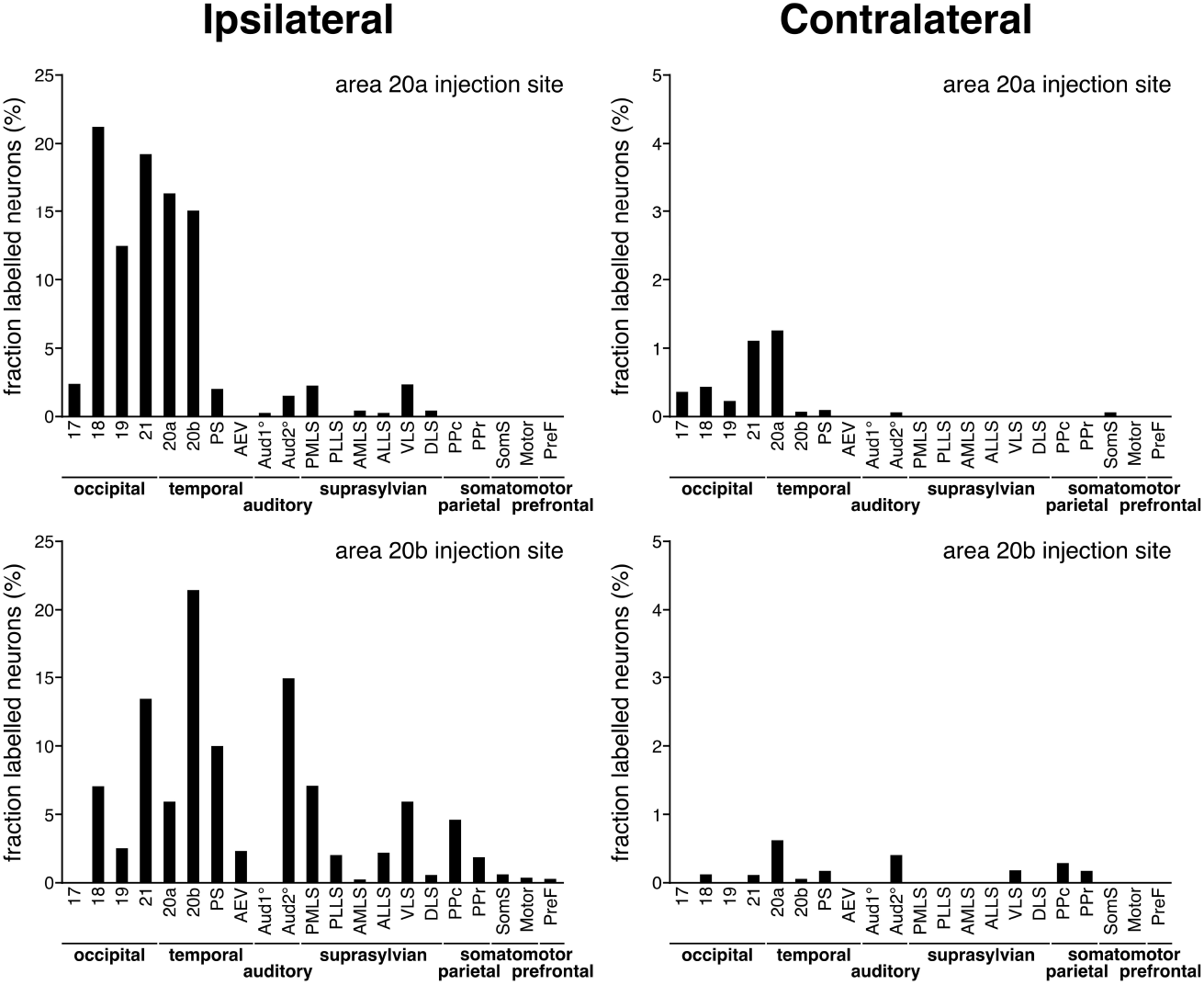
Graphs depicting the quantitative assessment of retrograde connectivity strength within and between cortical areas in the cerebral hemispheres ipsilateral (**left column**) and contralateral (**right column**) to the injection sites made in areas **20a** (**top two graphs**) and **20b** (**bottom two graphs**). The values are expressed in percentages, being the fraction of labelled neurons occurring in each cortical area. See list for abbreviations. Note that the a substantial proportion of retrogradely labelled cells are found within the cortical area injected, both ipsilaterally and contralaterally.

### Connectivity of area 20a

Injection of tracer into area **20a** resulted in both ipsilateral anterograde and retrograde label within the temporal, occipital and suprasylvian visual areas, temporal multisensory auditory/visual areas, with a small patch of anterograde label being observed in the ventral aspect of the pre-frontal cortex (Fig. 5). Extensive reciprocal connectivity was found within area **20a**, and also with areas **20b** and **PS** within the ventrolateral temporal visual areas. While all four occipital visual areas exhibited reciprocal connectivity to area **20a**, the strongest connectivity was observed in areas **18, 19** and **21**, area **17** having substantially weaker connections (Figs. 4, 5). While the extent of connectivity to the suprasylvian visual areas was weaker than that observed in the temporal and occipital visual areas, all six suprasylvian visual areas were connected with area **20a**, and all but area **DLS** showed reciprocal connectivity, with **DLS** only exhibiting retrograde connectivity to area **20a**. Again, less intense labelling was observed in the auditory and multisensory auditory/visual areas of the ectosylvian gyrus. Of the 8 different auditory, or auditory/visual multimodal, cortical areas defined in the ferret (Bizley, Nodal, Nelken, & King, 2005; Manger, Engler, Moll, & Engel, 2005), retrograde connections were observed in area **A1** and **AAF**, while areas **PSF** and **VP** were reciprocally connected to area **20a**, areas **AEV** and **PPF** received a minor anterograde projection, and areas **ADF** and **AVF** were not connected. A small patch of weak anterograde connectivity was found in the ventral region of the orbital gyrus, presumably representing prefrontal cortex (Fig. 5), and occasional cell bodies were identified in somatomotor (areas **M1**, **PV/SII**) and parietal (areas **PPr** and **PPc**) cortex. These qualitative impressions of the connectivity strength of the different cortical areas with area **20a** are supported by the quantification of the relative numbers of retrograde labeled neurons (N%) (Table 1, Fig. 4). The strongest ipsilateral retrograde connectivity was not observed within the area of the injection site (**20a**) but in area **18** (N% = 21.21%). This was followed by a decrease in N% in area **21** (N% = 19.26%), area **20a** (N% = 16.41%), area **20b** (N% = 15.10%), area **19** (N% =12.51), area **17** (N% = 2.39%), **VLS** (N% = 2.33%), **PMLS** (N% = 2.24%), and **PS** (N% = 2.07%). Thus, area **20a** appears to receive its most substantial feedforward input from the occipital visual areas.

**Figure 5:**
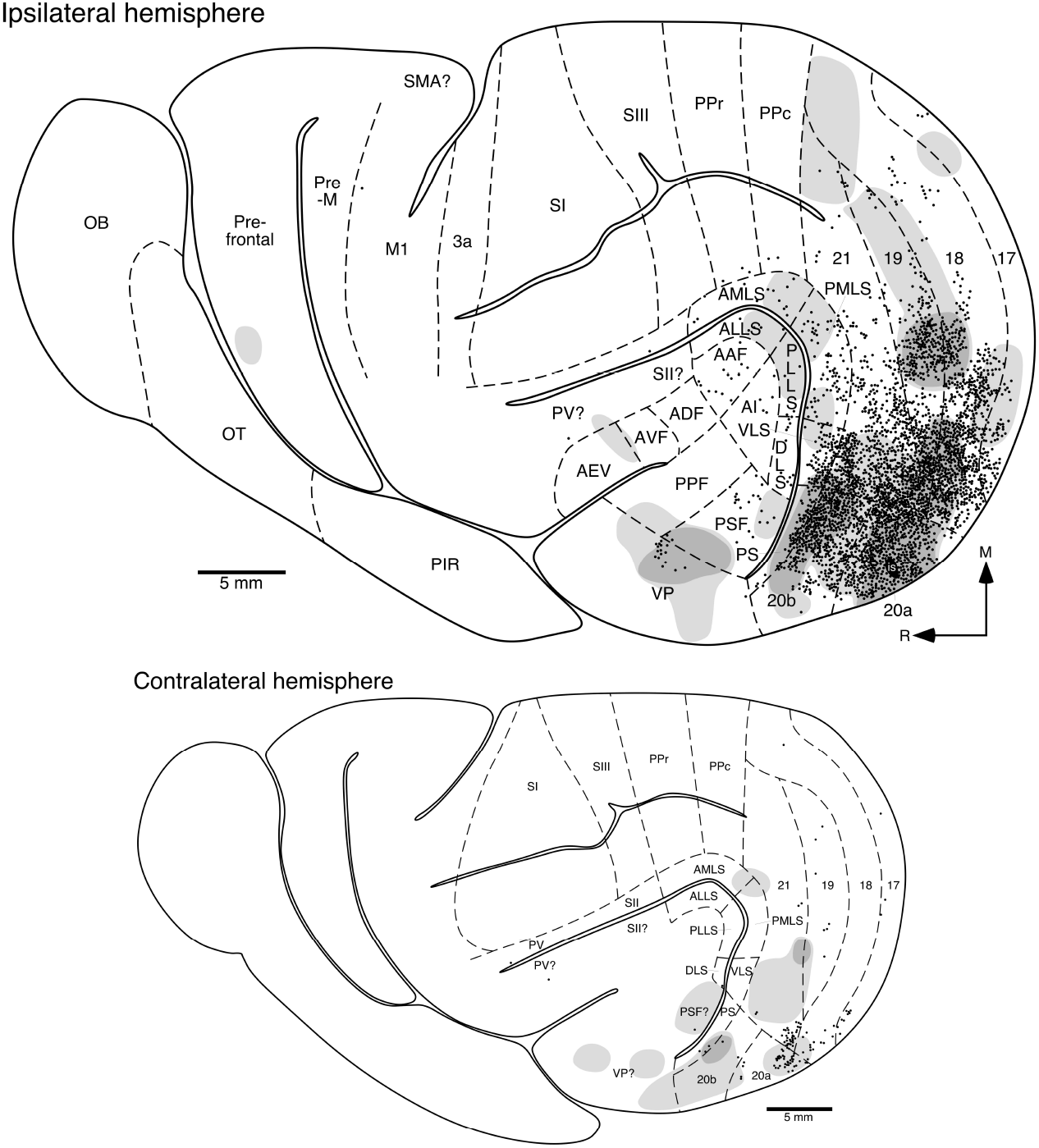
Location of retrogradely labelled cortical neurons (**filled circles**) and anterogradely labelled axons and axon terminals (dense labelling in the **darker grey shading**, light labelling in the **lighter grey shading**) following transport from the injection site (**is**) located in area **20a**. The upper larger image represents the distribution of cells and axons in the ipsilateral semi-flattened cerebral hemisphere, while the lower smaller image represents the semi-flattened cerebral hemisphere contralateral to the injection site. Note that ipsilaterally, extensive connectivity is seen through the occipital, suprasylvian and temporal visual and auditory areas, with a small patch of anterograde label found in the ventral aspect of the presumptive prefrontal cortex. The contralateral connectivity is much weaker and less widespread. Areal boundaries were demarcated using alternative sections stained for cytochrome oxidase and the boundaries represent approximations based on this stain and available maps of the ferret brain (Manger et al., 2002a,b, 2004, 2005; Manger, Engler, Moll, & Engel 2008; Manger, Restrepo, & Innocenti, 2010; Bizley et al., 2005; Homman-Ludiye et al., 2010). See list for abbreviations.

Contralateral connectivity following injection of tracer into area **20a** was far weaker than that observed ipsilaterally, and more restricted in terms of the areas covered (Fig. 5). The strongest reciprocal connectivity was observed in a region homotopic to the injection site. Retrogradely labeled cells were observed in the temporal visual areas **20a**, **20b** and **PS**, and in the occipital visual areas **17, 18, 19** and **21**, with the occasional cell observed in the contralateral somatosensory area PV and auditory area **PSF** (Table 1, Fig. 4). Anterograde projections were observed in the temporal visual areas **20a**, **20b** and **PS**, the occipital visual areas **18**, **19** and **21**, the suprasylvian visual areas **PMLS** and **AMLS**, and in the auditory areas **PSF** and **VP** (Fig. 5). Dense patches of anterograde label were only observed at the borders of areas **20b/PS** and **19/21**. Much of the contralateral connectivity was not reciprocal. These observations emphasize that the connectivity between the hemispheres is far weaker than that within the hemisphere, and that the contralateral connections are substantially less widespread than the ipsilateral connections.

Following injection of tracer into area **20a**, widespread, reciprocal connectivity was observed throughout all nuclei forming the visual dorsal thalamus, with the exception of the pulvinar and **P** lamina of the lateral geniculate nucleus, both of which only contained weak anterograde projections (Figs. 3a, 3d, 6). The reciprocal connectivity was the strongest throughout the MIN and the C laminae of the lateral geniculate nucleus, and the dorsal two thirds of **LP**. Weaker reciprocal connectivity was observed within the A lamina of the lateral geniculate nucleus, with minor anterograde projections to lamina **P** of the lateral geniculate nucleus and the pulvinar. Thus, area **20a** is strongly, and mostly reciprocally, connected with the majority of the nuclei of the visual portion of the dorsal thalamus of the ferret.

**Figure 6:**
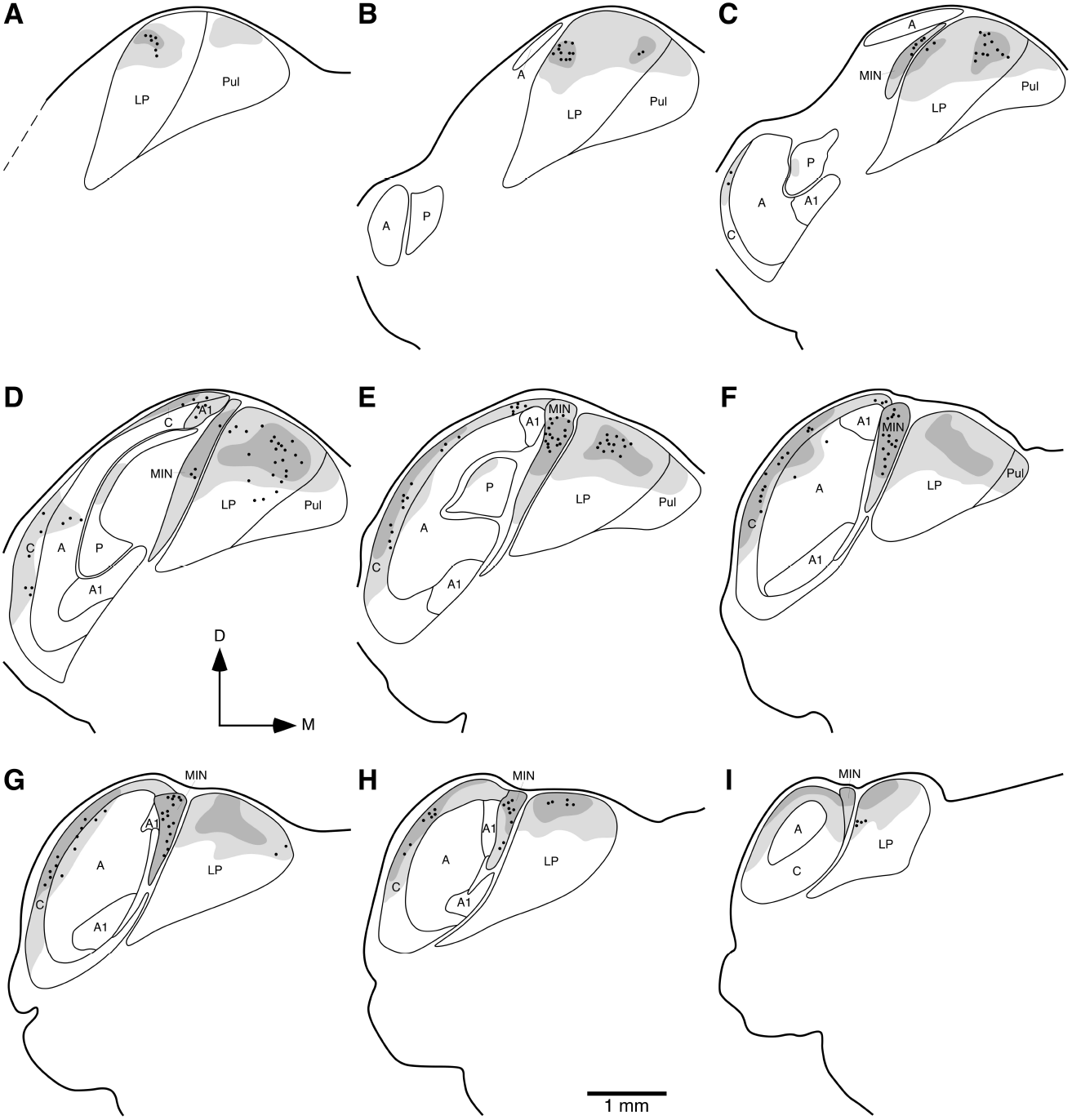
Diagrammatic reconstructions of the location of retrogradely labelled cells (**filled circles**) and anterogradely labelled axons and axon terminals (dense labelling in the **darker grey shading**, light labelling in the **lighter grey shading**) in the visual thalamus of the ferret following injection of tracer into the temporal visual area **20a**. a represents the most rostral coronal section, with i being the most caudal. Each section is approximately 400 μm apart. Note the dense, but restricted, label in the **C** lamina of the lateral geniculate nucleus and the strong connectivity to the **LP**. In all images dorsal (**D**) is to the top and medial (**M**) to the right. See list for abbreviations.

### Connectivity of Area 20b

Injections of tracer into area **20b** led to substantial and widespread ipsilateral reciprocal connectivity throughout much of the cerebral cortex (Fig. 7). The three posterior temporal visual areas, **20b**, **20a** and **PS**, all demonstrated strong reciprocal connectivity. Reciprocal connectivity was also observed in the occipital visual areas **18**, **19** and **21**, but no connections with area **17** were observed. Within the suprasylvian visual areas reciprocal connections were observed in **AMLS**, ALLS, **PMLS**, PLLS and **VLS**, while **DLS** only evinced retrogradely labeled cells. The parietal area **PPc** was reciprocally connected with area **20b**, but in **PPr** only retrogradely labeled cells were observed. Occasional retrogradely labeled cells were observed in the somatosensory areas **SII** and **SIII**, and the pre-motor region, while anterograde label was observed in the primary motor cortex (**M1**). A small patch of anterograde label and a few scattered retrogradely labeled cells were observed in the orbital gyrus, presumably prefrontal cortex (Fig. 7). Lastly, the auditory and auditory/visual multimodal areas of the ectosylvian gyrus exhibited extensive connectivity with area **20b** (Fig. 7). The auditory/visual areas **AEV** and **VP** showed extensive reciprocal connectivity to area **20b**, while weaker reciprocal connectivity was observed with the auditory areas **AVF** and **PPF**. The auditory visual area **PSF** exhibited a number of retrogradely labeled cells, presumably projecting to area **20b**. Quantitatively, the strongest ipsilateral retrograde connectivity was observed within area **20b** (N% = 21.42%) (Table 1, Fig. 4), with a decrease in connectivity strength in non-primary auditory cortex (N% = 14.93%), area **21** (N% = 13.40%), followed by **PS** (N% = 9.93%), **PMLS** (N% = 7.01%), area **18** (N% = 6.95%), area **20a** (N% = 5.95%), **VLS** (N% = 5.83%) and **PPc** (N% = 4.54%).

**Figure 7:**
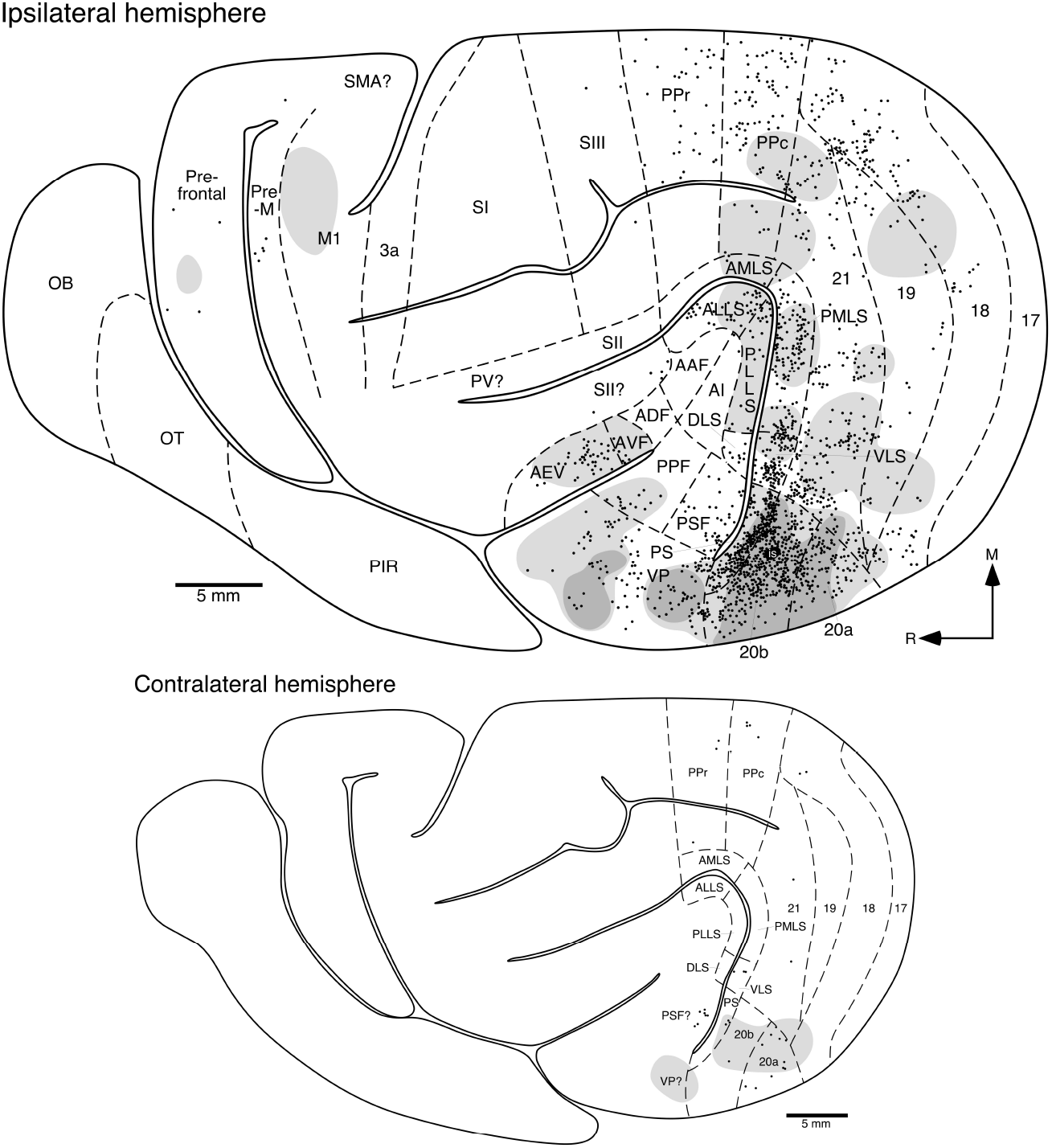
Location of retrogradely labelled cortical neurons (**filled circles**) and anterogradely labelled axons and axon terminals (dense labelling in the **darker grey shading**, light labelling in the **lighter grey shading**) following transport from the injection site (**is**) located in area **20b**. All conventions and abbreviations as provided in the legend to Fig. 5. Note the widespread and extensive ipsilateral connectivity throughout the occipital, suprasylvian, parietal and temporal, including auditory, areas. Also note the small number of retrogradely labelled cells and anterogradely labelled terminal fields in the motor and presumptive pre- frontal cortex. The contralateral connectivity, while not as strong as the ipsilateral, is restricted to the visual areas.

Contralateral connectivity following injection of tracer into area **20b** was extremely limited. Weak reciprocal connectivity was observed in the temporal visual areas **20b**, **20a** and **PS** (Fig. 7). Occasional retrogradely labeled cells were observed in areas **18**, **21**, **VLS**, **PPc**, **PPr** and **PSF**. Minor anterograde projections were observed in areas **18**, **21** and **VP**. Only two patches of anterograde labeling were observed in the contralateral hemisphere; however, this labeling was weak. The larger of the two patches extended from **PS** across to area **20a** to include the ventral boundary of area **18** and **21**. A second, patch extended across the area **20b/VP** border. This reiterates that the connectivity between the hemispheres is far weaker than that within the hemisphere, and that the contralateral connections are substantially less widespread than the ipsilateral connections following injections of tracer into area **20b**.

Following injection of tracer into area **20b**, widespread, mostly reciprocal, connectivity was observed throughout all nuclei forming the visual dorsal thalamus, with the exception of the **P** lamina of the lateral geniculate nucleus, which contained no labeling (Figs. 3b, 3c, 8). The most substantial reciprocal connectivity was observed within the **A** lamina of the lateral geniculate nucleus, the **LP** and pulvinar (Fig. 8). The anterograde labeling was evidenced as a dense core patches surrounded by a halo of weak to moderately strong terminal network densities. Minor reciprocal connectivity was observed in the C lamina of the lateral geniculate nucleus, and the **MIN**. Thus, area **20b** is strongly, and mostly reciprocally, connected with many of the nuclei of the visual portion of the dorsal thalamus of the ferret.

**Figure 8:**
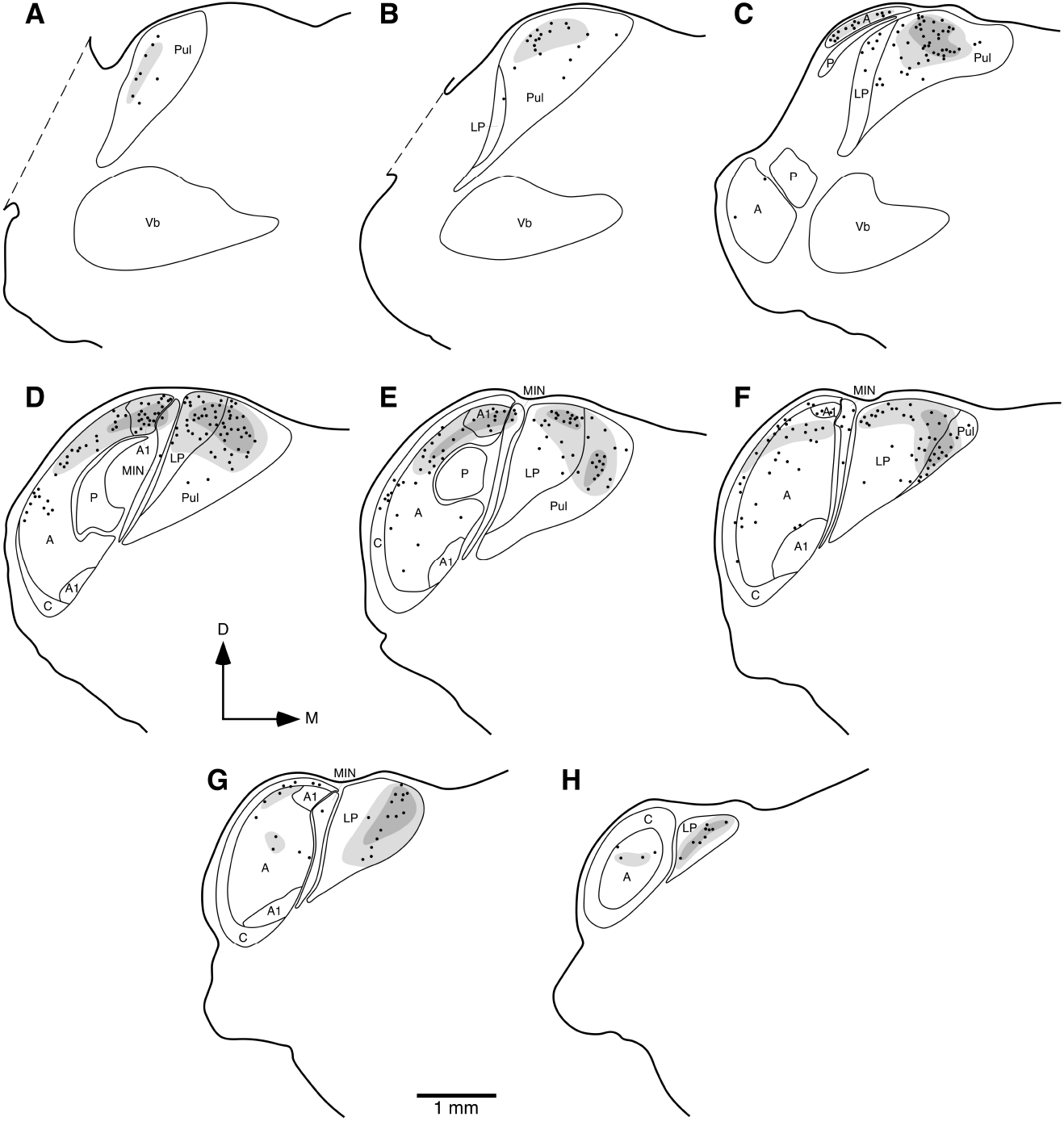
Diagrammatic reconstructions of the location of retrogradely labelled cells (**filled circles**) and anterogradely labelled axons and axon terminals (dense labelling in the **darker grey shading**, light labelling in the **lighter grey shading**) in the visual thalamus of the ferret following injection of tracer into the temporal visual area **20b**. **a** represents the most rostral coronal section, with **h** being the most caudal. Each section is approximately 400 μm apart. Note the dense connectivity with most regions of the visual thalamus, apart from that to the C lamina of the lateral geniculate nucleus and the **MIN**. Conventions and abbreviations as provided in the legend to Fig. 6 and abbreviation list.

## Discussion

The present study describes the ipsilateral and contralateral cortico-cortical and cortico-thalamic connectivity of the temporal visual areas **20a** and **20b** in the ferret using standard anatomical tract-tracing methods. These data will contribute to the Ferretome connectivity database (www.ferretome.org; Sukhinin, Engel, Manger, & Hilgetag, 2016), to facilitate cross-species analyses of brain connectomes and wiring principles of the brain. Areas **20a** and **20b** are not only well interconnected, but show numerous ipsilateral connections with occipital and temporal (visual and auditory) cortical areas, the latter providing the connectional foundation of previously observed physiological visual influences within the ferret auditory cortex (Bizley & King, 2009; Figs. 4, 9). However, the connectivity of area **20a** is denser than that of area **20b**, while the connectivity of area **20b** is more widespread than that of area **20a**. The contralateral hemispheres maintain a subset of the ipsilateral cortical projections (see Barbas, Hilgetag, Saha, Dermon, & Suski, 2005; Goulas, Uylings, & Hilgetag, 2017; Swanson, Hahn, & Sporns, 2017), and this connectivity was primarily anterograde and more limited for area **20b** than **20a** (Figs. 5, 7).

**Figure 9:**
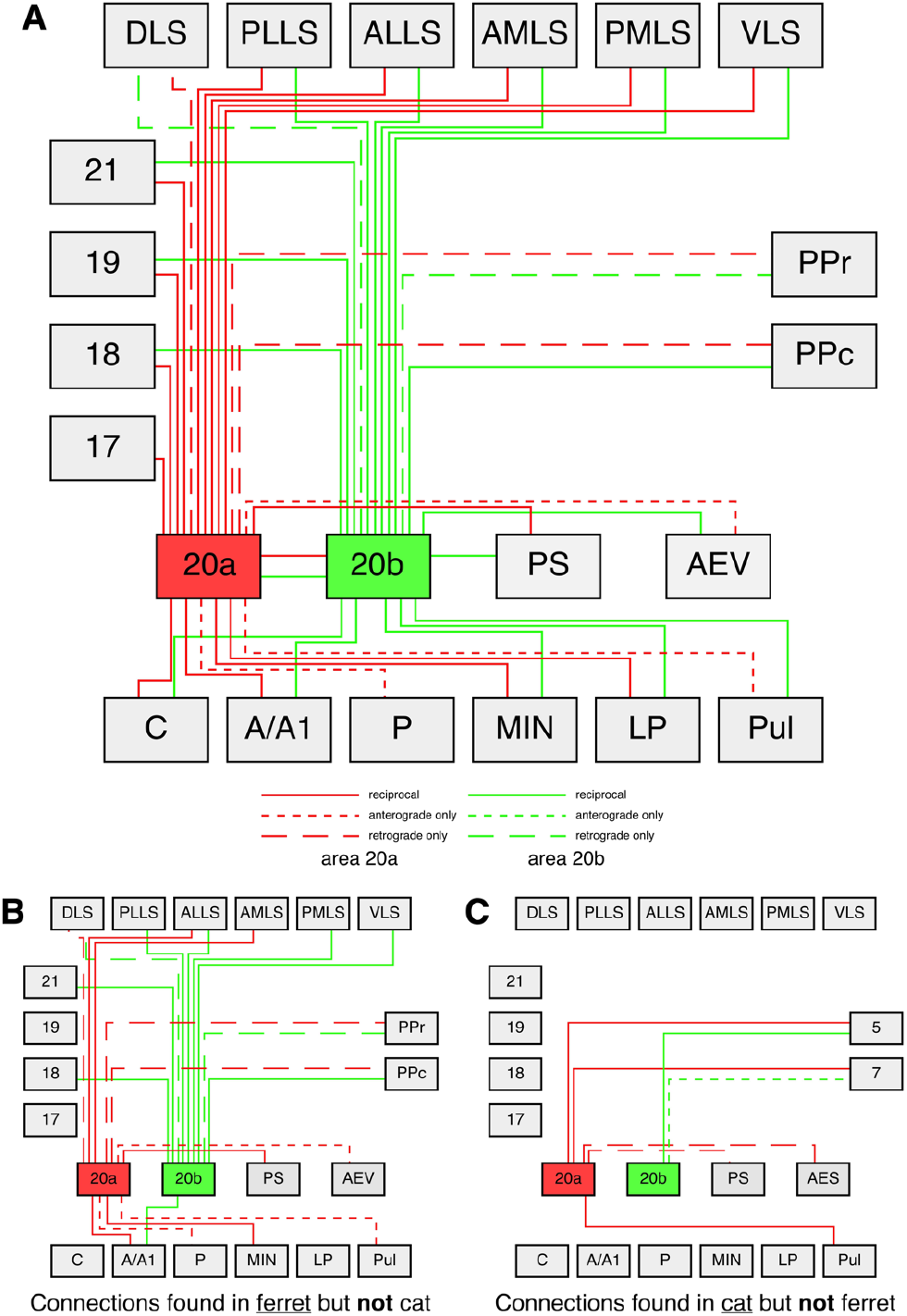
Wiring diagrams depicting the connectivity of areas **20a** and **20b** with each other, other visual cortical areas ipsilaterally, and the visual thalamus in the ferret (**a**), the connections of these areas observed in the ferret but not the cat (**b**), and the connections of these areas overs in the cat but not the ferret (**c**) (Heath & Jones, 1971; Symonds & Rosenquist, 1979, 1984; Symonds et al., 1981; Cavada & Reinoso-Suárez, 1983 Scannell et al., 1995, 1999). Each colour represents a specific cortical area (**20a** – red, **20b** – green), with solid lines representing reciprocal connections, short dashed lines anterograde connections only, and long dashed lines retrograde connections only. (**a**) Of the potential 120 connections (for each cortical area 20 potential reciprocal, anterograde only or retrograde only connections) of these two areas to the different visual cortical and thalamic regions, a total of 40 connections are found in the ferret. (**b**) In the ferret **21** connections not found in the cat were observed, while (**c**) in the cat 7 connections not found in the ferret were observed.

Comparisons to previous connectional studies of cat areas **20a** and **20b** (Fig. 9) showed numerous similarities and certain differences that may reveal a common pattern of connectivity of the temporal cortex for multisensory processing and integration within the cerebral cortex of carnivore and other mammals. As we noted earlier, it is difficult to propose direct temporal cortical area homologs of the ferret and cat with species with a more complexly organized temporal cortex such as the macaque monkey (see Payne, 1993; Manger et al., 2004). Thus we consider different regions, such as occipital, temporal, parietal, as a whole to elucidate whether global underlying connectivity patterns are present across species.

### Area 20a connectivity – ferret vs cat

Area **20a** in the ferret displays dense cortico-cortical ipsilateral connectivity, being reciprocally connected with temporal visual areas **20b** and **PS**, occipital visual areas **17**, **18**, **19** and **21**, and suprasylvian areas **ALLS**, **AMLS**, **PMLS** and **VLS**. It receives projections from areas **DLS**, **PPc** and **PPr**, and sends anterograde projections to area **AEV** (Figs. 5, 9). While cat area **20a** shares many of these connections, unlike the ferret it is reciprocally connected with areas 5 and 7 (ferret **PPr** and **PPc**), receives projections **PS** and **AES** (ferret **AEV**), and lacks connectivity with areas **ALLS**, **AMLS** and **DLS** observed for the ferret (Heath & Jones 1971; Symonds & Rosenquist 1984; Scannell, Blakemore, & Young, 1995). Area **20a** of the ferret displayed a more limited spread of contralateral connectivity in comparison to the ipsilateral connectivity, revealing sparse reciprocal callosal connections with areas **18**, **19**, **21**, **20a**, **20b** and **PS**, retrograde connections with area **17**, while sending projections to **AMLS** and **PMLS** (Figs. 5, 9). Area **20a** in the cat similarly evinced reduced callosal connectivity; however, the origins and targets of these projection neurons were dissimilar to those in the ferret, with reciprocal callosal connections only being observed in areas **20a** and **PMLS**, callosal projections received from areas **19** and **21** and callosal projections sent to areas **20b**, **DLS** and PLLS (Segraves & Rosenquist, 1982; Cavada & Reinoso-Suárez, 1983).

With regards to the connectivity of area **20a** with the visual nuclei of the dorsal thalamus, area **20a** in the ferret evinced connections with all the visual nuclei of the dorsal thalamus, with reciprocal connections to the **C**, **A** and **MIN** laminae of the lateral geniculate nucleus and the **LP**, and anterograde connectivity with the **P** lamina of the lateral geniculate nucleus and the pulvinar (Figs. 6, 9). In contrast, area **20a** in the cat displays limited connectivity with the visual nuclei of the dorsal thalamus, having reciprocal connections with the **C** lamina of the lateral geniculate nucleus, the **LP** and the pulvinar (Symonds, Rosenquist, Edwards, & Palmer, 1981; Scannell, Burns, Hilgetag, O’Neil, & Young, 1999). Thus, although area **20a** in the ferret and the cat share numerous similarities in connectivity patterns (Fig. 9), the connections with the anterior and dorsal lateral suprasylvian visual areas and with most of the lateral geniculate nucleus observed in the ferret are absent in the cat (Fig. 9). These differences reiterate our previous findings in the occipital and parietal visual areas of ferret as compared to the cat, where the higher order visual areas of the ferret are under a stronger influence of lateral geniculate input compared to the cat, potentially emphasizing differences in the extraction of visual information in the two species possibly related to life history and morphological differences (Dell, Innocenti, Hilgetag, & Manger, 2018a; Dell, Innocenti, Hilgetag, & Manger, 2018b).

### Area 20b connectivity – ferret vs cat

Area **20b** in the ferret, although not as heavily interconnected as area **20a**, exhibited expansive reciprocal connectivity with areas **18**,**19**, **21**, **20a**, **PS**, **AEV**, **ALLS**, **AMLS**, **PLLS**, **PMLS**, **VLS** and **PPc**, as well as retrograde connectivity with areas **DLS** and **PPr** (Fig. 7, 9). Area **20b** in the cat showed fewer connections than that of the ferret, revealing reciprocal connections only with areas **19**, **AMLS**, **20a**, **PS**, **AES** (ferret **AEV**) and 5 (ferret **PPr**), and projections to area 7 (ferret **PPc**) (Heath & Jones 1971; Symonds & Rosenquist 1984; Scannell et al., 1995). In contrast to area **20a**, area **20b** of the ferret had sparser contralateral connectivity, exhibiting reciprocal callosal connections with areas **18**, **21**, **20a**, **20b** and **PS**, while receiving callosal connections from areas **VLS**, **PPc** and **PPr** (Figs. 7, 9). Similarly, cat area **20b** maintained limited callosal connections, but the pattern of connectivity was different to that observed in the ferret, as reciprocal callosal connections were only identified with areas **21** and **20b**, while callosal projections were sent to areas **18** and **PMLS** and received from areas **19** and **20a** (Segraves & Rosenquist, 1982; Cavada & Reinoso-Suárez, 1983).

Area **20b** of the ferret exhibited reciprocal connections with all visual nuclei of the dorsal thalamus except for the **P** lamina of the lateral geniculate body where connections were absent (Figs. 8, 9). Area **20b** in the cat was similarly connected to all portions of the visual thalamic nuclei apart from the **A** and **P** lamina of the lateral geniculate nucleus which contained no connections (Symonds et al., 1981; Scannell et al., 1999). Although area **20b** in the ferret and the cat share numerous thalamic and cortical connections, numerous ipsilateral connections with the suprasylvian visual areas as well as callosal connections are absent in the cat. This finding could be indicative of specialized visual function and integration that would not serve the cat or it could simply be that these cortical connections are present in the ferret as area **20b** is in closer proximity with the suprasylvian visual areas and shares areal borders, particularly with **VLS**, whereas in the cat area **20b** is bounded by areas **17** and **20a** (Tusa & Palmer, 1980; Updyke, 1986; Manger et al., 2004).

### The role of the temporal visual cortex in the ferret

The current work demonstrates that areas **20a** and **20b** are not only well interconnected, but maintain connections with numerous visual and secondary auditory areas (Fig. 4; Table 1). The connections with auditory cortex supplements the finding by Bizley & King (2009) that visual stimulation of area 20 in the ferret modulates activity of neurons in the auditory cortex. Thus, it is plausible that area 20 in the ferret is a possible site for the integration of auditory-visual information; however, the explicit integration of multisensory stimuli would have to be tested by presenting the unisensory stimuli simultaneously and comparing this to instances where the stimuli are presented in isolation (see Beauchamp, 2005). Interestingly, our findings show that area **20a** has more dense connections with the occipital cortex while area **20b** is more densely connected with the secondary auditory cortex (Fig. 4, 5, 7; Table 1). This can be expected, as areas **20a** and **20b** are located between the auditory and occipital cortices, with **20a** being in closer proximity to the occipital cortex and **20b** to the auditory cortex respectively (see Tusa & Palmer, 1980; Updyke, 1986; Wallace, Meredith, & Stein, 1992; Manger et al., 2004; Fig. 5). Furthermore, the location of areas **20a** and **20b** between two larger unimodal cortical regions, visual and auditory, could further explain the multisensory aspects of neural information processing in areas **20a** and **20b**, as cortical regional border zones tend to be biased towards multisensory representation (Wallace, Ramachandran, & Stein, 2004; Ghazanfar & Schroeder, 2006; Dahl, Logothetis, & Kayser, 2009). Cortical multisensory integration requires a site where different information modalities can converge (Wallace et al., 1992), and areas **20a** and **20b** provide such a site for the integration of visual and auditory information. In addition, area **20b** is more strongly connected with the posterior parietal cortex than area **20a** (Fig. 4, 6; Table 1), thus allowing area **20b** to play a role in dorsal stream processing (Goodale & Milner, 1992; Goodale, Kroliczak, & Westwood, 2005). Thus, although the temporal visual cortex is a site for processing ventral stream visual information (object perception), it may also function, in the ferret, as a site that integrates multisensory stimuli to provide a comprehensive scene in which objects can be unambiguously perceived.

### Interaction of the dorsal and ventral streams

In order for aspects of a visual image to be fully extracted by the brain, the dorsal and ventral streams need to interact, and the prefrontal cortex is involved in this process (Miller, 2000; Itti & Koch, 2001). The posterior parietal cortex has been identified as an integral component of the visual dorsal stream, maintaining direct connections with the prefrontal cortex, as well as numerous direct and indirect connections with the occipital cortex. This has been illustrated in numerous animal studies (Baizer, Ungerleider, & Desimone, 1991; Remple, Reed, Stepniewska, Lyon, & Kaas, 2007; Kaas & Stepniewska, 2016) as well as in the ferret (Dell et al., 2018a,b). Similarly, the visual temporal cortex is integral to visual ventral stream information processing, thus, direct connections with the prefrontal cortex are anticipated (Goodale & Milner, 1992; Webster et al., 1994; Cloutman, 2013). Our study of the visual temporal cortex in the ferret revealed that both areas 20a and 20b are connected with the prefrontal cortex (likely the orbitofrontal cortex), but these connections were very sparse (Figs. 4, 5, 7; Table 1).

Given that integration of the ventral and dorsal processing streams is required for full visual perception, and that direct connectivity of the temporal visual areas with the prefrontal cortex is weak in the ferret, how does information from the ventral processing stream reach the prefrontal cortex to allow for integration? As area **20b** in the ferret is well connected with the posterior parietal cortex (Fig. 4, 7; Table 1), we can speculate that information from the ventral processing stream passes from area **20b** to the posterior parietal cortex and then to the prefrontal cortex for sensory integration. Thus, the posterior parietal cortex may be the primary site for dorsal and ventral stream interaction in the ferret. This speculation is congruent with findings that show that cortical areas of the monkey inferior temporal lobe (areas **TEO** and **TE**), responsible for ventral stream processing, are connected to the posterior parietal cortex (Webster et al., 1994). Similarly, cat temporal visual cortex maintains connections with posterior parietal areas 5 and 7 (Symonds & Rosenquist, 1984).

Moreover, results from our previous study (Dell et al., 2018a), indicated that area **PMLS** is directly connected to occipital visual areas **17**, **18**, **19** and **21**, implicating it as a hub that may serve to connect the temporal visual areas to the parietal cortex for dorsal stream processing. The current study shows that areas **20a** and **20b** are well connected with **PMLS** (Fig. 4,5,7,9; Table 1), a feature also observed in the cat (Symonds & Rosenquist, 1979). Thus, information from the ventral processing stream can be sent to area **PMLS** and the posterior parietal cortex, regions identified for their explicit involvement in dorsal stream processing (Goodale & Milner, 1992; Goodale, 1998; Goodale et al., 2005; Cloutman, 2013). Hence, we can speculate that **PMLS** and the posterior parietal cortex (**PPc** and **PPr** in the ferret) may be two intermediary integration sites, or hubs, where the dorsal and ventral streams interact prior to the visual information being sent to the frontal cortex.

This potential connectional network is valid for **PMLS** and areas **5** and **7** in the cat (homologues of **PPr** and **PPC** in the ferret); however, it is more challenging to apply a similar network framework to species such as the macaque monkey as the homologies of cortical areas between cat or ferret and macaque monkey are unclear, and the parietal and temporal cortical regions have undergone expansion and increased parcellation in the macaque (see Baylis, Rolls, & Leonard, 1987; Payne, 1993; Manger, Masiello, & Innocenti, 2002b; Manger et al., 2004; Kaas and Stepniewska; 2016). As alluded to earlier, macaque monkey inferior temporal cortex (ventral stream processing) is strongly connected with the posterior parietal cortex as well as the prefrontal cortex (Webster et al., 1994), with area TE of the macaque temporal cortex being well connected with area **MT** (Webster et al., 1994). Moreover, if one considers that area **MT** in the macaque monkey is analogous to area **PMLS** in the carnivores (Lappe & Rauschecker 1995; Cantone, Xiao, & Levitt, 2006), it appears that the dorsal and ventral processing streams interact in the posterior parietal cortex as well in the central temporal region (suprasylvian area **PMLS** for ferret and cat, and **MT** for macaque monkey) enroute to integration in the prefrontal cortex. Such a stepwise network interaction may ease the processing load on the prefrontal cortex and minimize the wiring cost of the brain (see Laughlin & Sejnowski, 2003; Bullmore & Sporns, 2012). Further research needs to be done in other species to identify and confirm additional sites of possible dorsal and ventral stream interaction. Nevertheless, it appears that the ferret possesses, at a minimum, the basic connectivity network required for dorsal and ventral stream integration, and in a manner not dissimilar to that observed in the primates.

## Acknowledgements

The authors wish to thank Mrs. Sonata Valentiniene for her consistently high-quality histological preparations.

## Role of Authors

PRM and GI designed the study and undertook the experimental aspects of the study. LAD, CCH and PRM analyzed the material and LAD wrote the first draft of the paper, which was subsequently edited by GMI, CCH and PRM. All authors had full access to all of the data in the study and take responsibility for the integrity of the data and the accuracy of the data analysis.

